# Xenotransplantation of porcine progenitor cells in an epileptic California sea lion (*Zalophus californianus*)

**DOI:** 10.1101/2021.07.30.454497

**Authors:** Claire A. Simeone, John P. Andrews, Shawn P. Johnson, Mariana Casalia, Ryan Kochanski, Edward F. Chang, Dianne Cameron, Sophie Dennison, Ben Inglis, Gregory Scott, Kris Kruse-Elliott, F. Fabian Okonski, Eric Calvo, Kelly Goulet, Dawn Robles, Ashley Griffin-Stence, Erin Kuiper, Laura Krasovec, Cara L. Field, Vanessa F. Hoard, Scott C. Baraban

## Abstract

**Background:** Domoic acid (DA) is a naturally occurring neurotoxin harmful to marine animals and humans. California sea lions exposed to DA in prey during algal blooms along the Pacific coast exhibit significant neurological symptoms, including epilepsy with hippocampal atrophy.

**Observations:** Here we describe a xenotransplantation procedure to deliver interneuron progenitor cells into the damaged hippocampus of an epileptic sea lion with suspected DA toxicosis. The sea lion has had no evidence of seizures following the procedure, and clinical measures of well-being including weight and feeding habits have stabilized.

**Lessons:** These preliminary results suggest xenotransplantation has improved the quality-of-life (QOL) for this animal and holds tremendous therapeutic promise.

## Introduction

Domoic acid (DA) is a neurotoxin produced by *Pseudo-nitzschia* algae^1^. DA toxicosis of California sea lions (*Zalophus californianus*) that consume the toxinis now widespread and a common cause of morbidity and mortality^2^. Unusually aggressive behaviors, vomiting, inappetence, marked lethargy, stranding in atypical locations, ataxia, seizures, coma and increased mortality are observed in DA-intoxicated sea lions rescued for rehabilitation in California^3–6^. Many of these animals exhibit an acquired form of epilepsy (i.e., spontaneous recurrent seizures, hippocampal atrophy, mossy fiber sprouting and interneuron cell loss) resembling human mesial temporal lobe epilepsy (TLE) ^7^. Similar to many human TLE patients, treatments to control seizures in DA-intoxicated sea lions are urgently needed. Available options for wild animals only address acute toxicosis while in rehabilitation, and DA-exposed sea lions placed in long-term care can develop progressive disease despite administration of antiepileptic drugs (AEDs)^6^.

A one-time cell transplantation therapy, using GABAergic interneurons that efficiently integrate in local epileptic circuits, and function similar to endogenous inhibitory neurons, with the potential to correct co-morbid behavioral deficits, would offer an alternative. GABAergic progenitor cells derived from embryonic medial ganglionic eminence (MGE) integrate into host circuits following transplantation, where they make functional inhibitory synapses^8^, efficiently suppress seizures, and correct co-morbid behavioral deficits^9,10^. These observations are based on harvesting embryonic murine MGE progenitors and rodent acquired epilepsy models. Building on these studies, we recently established a protocol to harvest porcine MGE progenitors as a tissue source for xenotransplantation^11^. Progenitor cell xenotransplantation into marine mammals, specifically epileptic sea lions, has never been attempted but, if successful, could provide a route to disease-modifying therapies.

## Illustrative Case

We report a pilot case of intra-hippocampal porcine progenitor cell xenotransplantation in a California sea lion with refractory epilepsy and magnetic resonance imaging (MRI) characteristics of unilateral hippocampal atrophy.

### Epilepsy

The observed course of epilepsy in the subject is as follows: subject first stranded on the coast of San Luis Obispo county, CA in November 2017, was admitted to a rehabilitation center for lethargy and disorientation, but recovered rapidly and was released two weeks later. Over the next two months, the sea lion was rescued twice more. Veterinarians determined the sea lion was unlikely to survive in the wild due to habituation. He was deemed non-releasable by NOAA Fisheries and transferred to a facility for permanent care. A convulsive seizure with impaired consciousness was observed in April 2018. A second seizure was observed in February 2019 and phenobarbital treatment was initiated as a first line AED. MRI obtained in January 2018 showed no structural abnormalities, though a second MRI obtained in October 2018 showed evidence of unilateral hippocampal atrophy (Figure 1A, top panel). Over the next 14 months, three additional convulsive seizures were observed (Figure 1B, top panel) that were associated with periods of prolonged anorexia and reduced feeding. (Figure 1B, middle and bottom panel). Diazepam was added as a second AED in April 2020.

**Figure 1:**
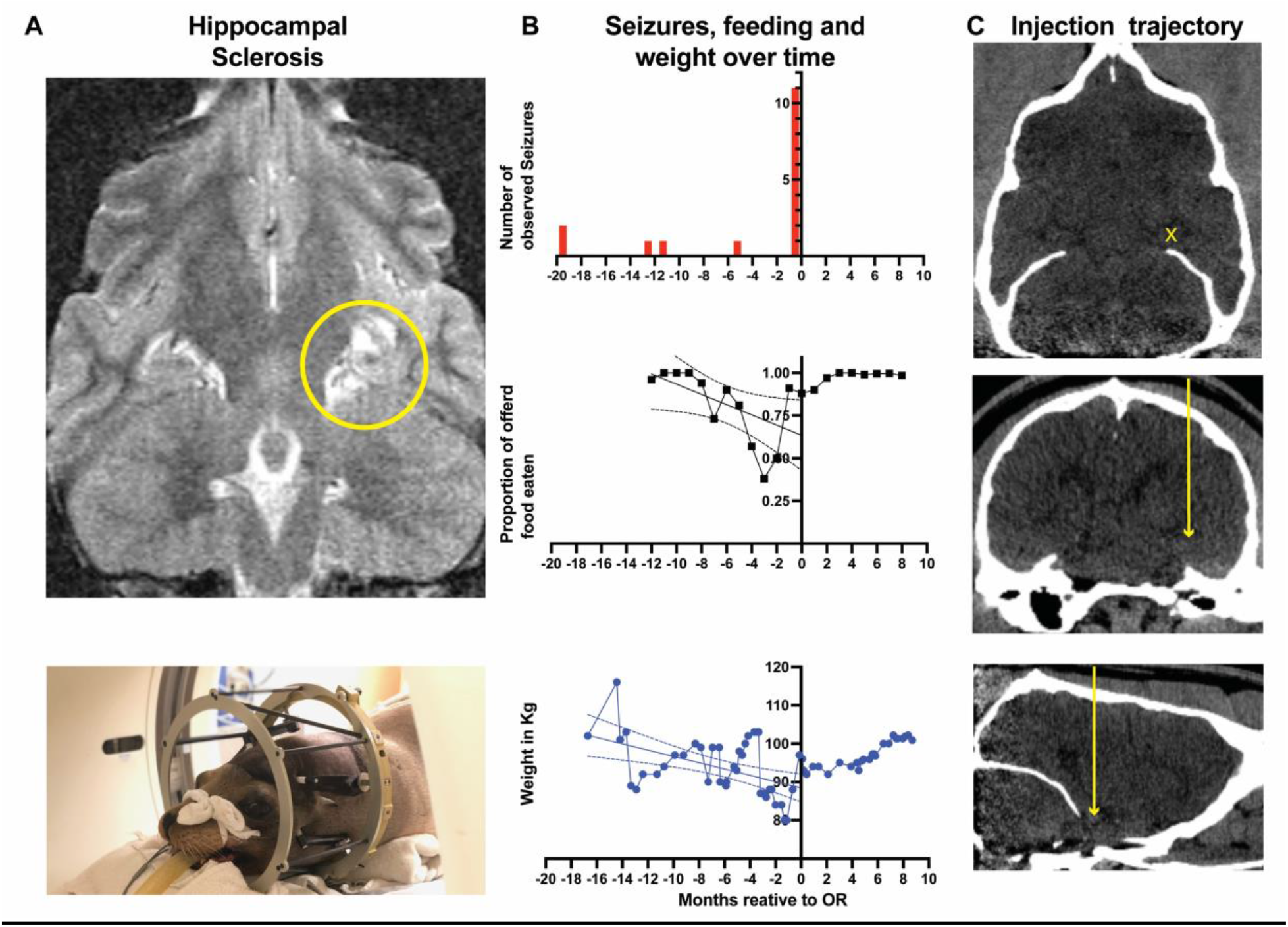
**A)** Transverse (axial) slice of a T2-weighted MRI of the sea lion subject (top panel). The yellow circle outlines the left hippocampus showing relative atrophy compared to contralateral hippocampus, as evidenced by increased T2-bright CSF spaces of the temporal horn of the lateral ventricle. Bottom panel shows the subject fitted with a stereotactic frame, intubated for a pre-procedural CT-scan (**B**) Observed seizures (top panel), proportion of food eaten as a fraction of total food offered (middle) and average monthly weight in kilograms (Kg) over time (in months), relative to date of the procedure (at months = 0). Trend lines for variables preintervention are overlaid onto middle and bottom panels are linear regressions with 95% confidence intervals of pre-intervention time-points (i.e. not including points after months = 0). R^2^ of proportion of food eating = 0.32; R^2^ of weight = 0.23. (**C**) CT scan immediately prior to procedure, showing transverse (top), dorsal (middle), and sagittal (bottom) slices. Yellow crosshairs delineate hippocampus target (top panel); yellow arrows delineate injection needle trajectory.

In early 2020, complete anorexia (1-3 days per month progressing to 6-17 days per month) was noted. Food consumption and monthly average weight show a clear declining trend (Figure 1 B). Appetite stimulants (mirtazapine and capromorelin) were offered with minimal observed therapeutic effect; ~25% body weight loss (from 103 to 79 kg) was noted during this period. Progression in anorexia coincided with eleven more observed seizures over a 5-day period in September 2020, despite therapeutic levels of phenobarbital.

### MGE Xenograft

With an inability to control seizures, increasing episodes of anorexia, and declining body weight, euthanasia was considered. As an alternative, a scientific-therapeutic collaboration led to a single-subject xenotransplantation trial modified to accommodate a large marine mammal. Under anesthesia, a computed tomography (CT) scan was performed to map surgical trajectory to the left hippocampus, and the head fixed in place for stereotactic implantation. Progenitor cells harvested from porcine embryos were prepared ^11^. 50,000 cells/site of MGE progenitors were targeted for locations along a single tract trajectory to the left hippocampus (Figure 1C). At 9post-transplantation no seizures have been observed (Figure 1B, top panel), appetite and weight have stabilized (Figure 1B, middle and bottom panels), and behaviors appear subjectively improved. The animal has continued post-surgery AED treatment (phenobarbital and diazepam) to reduce the risk of withdrawal seizures^12,13^.

## Materials and Methods

### Subject

The subject was a 8-year-old male California sea lion (*Zalophus californianus*) clinically diagnosed with epilepsy from earlier suspected DA toxicosis based on observed seizure-like activity, prolonged anorexia, radiologic evidence of unilateral hippocampal atrophy and absence of infectious disease agents.

### Pharmacological treatments

Immunosuppression consisted of cyclosporine (7 mg/kg SID for 7 days, then 3 mg/kg BID for 2 months, then 2 mg/kg SID) and dexamethasone (0.17 mg/kg SID for 2 weeks, tapered to 0.1 mg/kg SID) that were administered immediately prior to the procedure and for six months afterwards to prevent MGE cell rejection. Cyclosporine serum levels were monitored (Clinical Pharmacology Laboratory, Auburn University) with 24-hour trough levels of 7 mg/kg SID of 183-280 ng/mL. Prophylactic doses of antibiotics (ceftiofur, 6.6 mg/kg) were administered during the perioperative period and for one week postoperatively. Long-acting buprenorphine SR (0.2 mg/kg q72h) was used for ten days for analgesia. AED treatment consists of phenobarbital (1.6 mg/kg SID) and diazepam (0.05 mg/kg BID).

### Xenotransplantation procedure

The subject was anesthetized, intubated and maintained on isoflurane and oxygen. Freezethawed porcine embryonic MGE progenitor cells were prepared and >90% viability confirmed on site, as described^11^. A custom-made 10 cm long Hamilton needle (1.8 mm internal diameter; two outer sections 5 cm length, 23G followed by 5cm 32G blunt) was back-loaded with porcine progenitor cells. An entry point was chosen rostral and lateral to occipital eminence, from which a trajectory orthogonal to skull was used to enter hippocampus. The sea lion was positioned in sternal recumbency, prepped and draped in sterile fashion. A stab incision was made in the skin and twist drill was used to make a small burr hole for the injection needle. The MGE-loaded Hamilton needle was inserted slowly to a pre-measured depth based on pre-operative CT scan and progenitor cells injected at a rate of 5 nl/sec in 3 locations along the trajectory: once at initial target depth, followed by a withdraw of approximately 5 mm, repetition of injection procedure, then withdrawing 5 mm and injecting once more. Following cell delivery, injection needle was slowly withdrawn, and skin closed with an absorbable suture. The sea lion was maintained in a dry pen for three days postoperatively.

## Discussion

### Observations

Our compassionate-use trial in a thrice-stranded, severely anorexic, presumed DAexposed sea lion with multiple documented episodes of convulsive seizures attributed to a hippocampal lesion, suggests porcine GABA progenitors are particularly promising candidates for xenotransplantation-based cell therapy. The rationale for attempting this life-saving therapy is based upon studies regarding origins and properties of inhibitory interneurons from embryonic MGE^14^. Control of seizure activity and rescue of co-morbid behavioral deficits associated with acquired epilepsies was initially described in mouse models using murine embryonic MGE progenitor cell donors transplanted into a chemoconvulsant damaged hippocampus^9,10^. Murine MGE-derived cells migrate widely and differentiate to functional GABA-expressing interneurons capable of enhancing synaptic inhibition in these mice. Similar to AEDs that enhance GABA-mediated inhibition (i.e., benzodiazepines) therapeutic benefit of MGE transplantation is likely associated with addition of inhibitory interneurons into a hyper-excitable hippocampal network. To translate this strategy to larger mammals we adapted a protocol using fetal pigs, a well-established source for xenotransplantation tissue ^15^. Recent demonstration that porcine embryonic MGE progenitors migrate and differentiate into GABA-expressing interneurons in a manner similar to that described for murine embryonic MGE progenitors used for cell transplantation therapy^11^, established this porcine cell source as a viable candidate for treating larger animals. Because California sea lions with DA toxicosis represent a wellcharacterized example of acquired epilepsy in wildlife^5^, and prior studies describe unilateral hippocampal atrophy with loss of inhibitory hippocampal interneurons in some of these animals^7,16^ it is not entirely surprising that successful MGE-based treatment of a sea lion mimics prior mouse studies.

Here we followed immunosuppression protocols established for transplantation of embryonic dopaminergic cells in Parkinson disease patients. As cell survival was noted 24 years after these procedures ^17^ this risk may not be a limiting factor, and potential to generate pathogen-free pigs as tissue donors could further mitigate concerns. A limitation of the current procedure was the inability for image verification during, or after, intra-hippocampal delivery of MGE progenitors. Real-time MRI-guided xenotransplantation may be possible with co-injection of a contrast agent ^18^ to confirm anatomic localization and cell delivery at the injection site. Long term monitoring of MGE-derived cells may be more difficult as they disperse and migrate in host brain thus requiring single-cell resolution neuro-imaging techniques.

### Lessons

Although pre-procedure monitoring of the subject’s behavior and epilepsy was limited, and these studies describe a relatively short survival period, improvement in QOL measures and absence of observable convulsive seizures for over nine months offers a cautious enthusiasm. DA toxicosis is the most common cause of neurological abnormalities in stranded California sea lions^19^ and expected climate-driven increases in HABs will continue to result in hundreds of sea lions (and other marine mammals) with DA toxicosis annually^1^ that could potentially benefit from such therapy.

## Acknowledgements

Barbie Halaska, Abby McClain, Frances Gulland from the Marine Mammal Center.

